# Laser Speckle Orthogonal Contrast Imaging Calibration by Replicating Red Blood Cells Scattering Statistics with a Moving Reference Diffuser

**DOI:** 10.64898/2025.12.03.692151

**Authors:** Xavier Orlik, Aurélien Plyer, Elise Colin

## Abstract

Recent studies have proposed improving Laser Speckle Contrast Imaging (LSCI) by using polarimetric filtering to isolate multiply scattered photons from moving red blood cells (RBCs), an approach referred to as Laser Speckle Orthogonal Contrast Imaging (LSOCI). This reliance on multiple scattering enables the development of a calibration method based on a moving reference sample, chosen to generate dynamic circular Gaussian speckle fields that replicate the statistical properties of RBC scattering in both intensity and the distribution of polarization states. Assuming that multiply scattered photons from both RBCs and the reference sample exhibit a homogeneous distribution of polarization states over the Poincaré sphere, the proposed calibration links *in vivo* speckle contrast reduction in a bijective manner to an equivalent speed of the reference sample. We demonstrate that this equivalent-velocity metric yields consistent *in vivo* measurements across distinct instruments despite the use of different laser spectral widths, thereby providing a standardized and transferable means to quantify microcirculation activity.

## 1 Introduction

Dynamic speckle imaging (DSI)^1– 3^has emerged as a remarkably attractive optical technique for probing biological dynamics, particularly in medical applications involving tissue microcirculation. Its technological simplicity, low cost, and non-invasive nature make it suitable for real-time, wide-field imaging of processes such as blood flow or cellular motility. Among the various DSI implementations, Laser Speckle Contrast Imaging (LSCI) and its polarimetric extension, Laser Speckle Orthogonal Contrast Imaging (LSOCI), have shown strong potential for clinical translation by enabling high-resolution, contrast-agent-free visualization of functional tissue perfusion. The principle of LSOCI is as follows: by illuminating a region of interest on the body with fully polarized laser light and detecting only the orthogonally polarized component of the backscattered field, a stable, high-resolution subsurface microcirculation network can be revealed with remarkable clarity. In fact, the portion of incident light that undergoes single scattering, primarily due to surface interactions, retains the incident polarization and is filtered out by the detection system. In contrast, photons penetrating deeper layers experience multiple scattering events, leading to spatial depolarization that spans the entire Poincaré sphere.^4^ By selectively filtering out the incident polarization, only photons that have undergone multiple scattering in subsurface layers are detected. Thus, the backscattered field can be decomposed into two main contributions : the surface contribution, which is filtered out, and the volume contribution, which encapsulates the temporal dynamics of multiple scattering processes.

Another critical aspect of LSOCI is the use of an appropriate near-infrared wavelength, which achieves optimal penetration depth while being particularly suited for multiple scattering interactions with red blood cells. Depths of imaging up to 3 mm have already been reported,^5^ and even larger penetration depths may be achievable depending on the anatomical region.

However, to fully harness its potential, it is essential to provide quantitative microcirculation measurements alongside images. Indeed, quantifying local RBC movement or agitation can be particularly valuable for applications such as detecting and monitoring therapies of skin tumors, assessing deep burns, evaluating inflammatory conditions, and more generally investigating various pathologies affecting the microcirculation network. Thus, in the general framework of DSI, several calibration methods have been proposed to measure absolute blood flow.^6–10^ However, they seem better suited to research settings than to clinical practice because of their rather complex implementation. Moreover, none of them takes into account the spectral width of the laser used, which may hinder direct inter-comparison of the measurements reported across these studies. Only the work of D. Postnov et al.^11^ proposed some experimentations and numerical simulations to quantify the effect of the spectral width of the laser on the speckle contrast, upon which measurements are based.

In the specific context of LSOCI, the signal relies on multiple scattering, which complicates establishing a direct relationship between temporal speckle contrast variation and red blood cell absolute speed. In fact, detected photons undergo multiple scattering within an elementary volume where RBCs move potentially in three-dimensional (3D) patterns. Although speckle transverse and longitudinal correlation lengths differ, determining the vectorial speed contributions of RBCs that cause contrast reduction remains practically intractable.Recognizing these inherent limitations, we adopt a different viewpoint: instead of attempting to recover an absolute flow velocity, we propose to measure a global 3D agitation metric of RBCs. This perspective is well aligned with the dense, volumetric architecture of the microvascular network, where the local motion of multiple RBCs jointly contributes to speckle decorrelation.

In this context, we introduce a new calibration method for LSOCI that delivers reproducible quantitative results. This calibration accounts for the spectral width of the laser, the detection optics and the detector characteristics. Thus, it enables effective comparisons across different LSOCI instruments and facilitates standardized microcirculation activity assessments. To enable this calibration, we employ a dedicated moving reference sample, for which the detected temporal contrast variation as a function of speed is mapped onto the *in vivo* contrast changes induced by the motion of RBCs.

The next section of the paper provides a detailed description of the calibration procedure, including the rationale behind the choice of the moving reference sample. Section 3 presents the experimental results: it first reports the calibration of three dynamic speckle imaging systems using lasers with spectral widths of 0.001, 0.1, and 0.44 nm, and then compares *in vivo* microcirculation measurements acquired on the same finger-wound region with two independently calibrated LSOCI systems. Section 4 offers a discussion of these findings. Finally, the concluding section summarizes the main outcomes of the study and outlines the prospects opened by this calibration approach.

## 2 Calibration procedure

### 2.1 Choice and requirements of the multiple-scattering reference sample

As a first step, we need to emphasize that any dynamic speckle measurement is intrinsically a measure of the decrease in speckle contrast from a reference speckle contrast value. The latter value represents the maximal speckle contrast the imaging system can exhibit for a given integration time. It is dependent on the optical aperture, on the detector, and on the spectral width of the laser used. We define this value as the spatial speckle contrast detected on a rough static reference sample that backscatters the illuminating laser light. The roughness of this sample (standard deviation of the height distribution) must reach at least one third of the wavelength, and the illumination conditions must ensure that at least a dozen roughness cells (described by the correlation lengths of the surface height distribution) are illuminated.^12^ These conditions enable the generation of a circular Gaussian speckle-backscattered field. If, moreover, the sample exhibits a high reflectance at the wavelength used, multiple scattering will occur and provoke spatial depolarization, thus filling the entire Poincaré sphere.^4^ This reference sample will hereafter be referred to as an MS-CGS sample (for Multiple Scattering–Circular Gaussian Speckle sample), and accordingly, the proposed calibration procedure will be termed the MS-CGS calibration method. Such a sample is expected to mimic, from the point of view of statistical distribution, both the intensity and the polarization states of the backscattered field from RBCs. To support this hypothesis, note that RBCs, which are micrometer-sized, exhibit surface topography variations larger than the near-infrared wavelength used here. Moreover, the very large number of RBC-scattering contributions involved ensures that the Gaussian regime is reached. Any reference material satisfying these criteria qualifies as a *Multiple-Scattering Circular-Gaussian Speckle* (MS-CGS) sample. Although several calibrated diffusers could be used, a simple sheet of white Canson paper has been shown experimentally to satisfy all requirements.^13^

It is precisely this universality of the statistics describing a circular Gaussian speckle generated by multiple scattering that allows LSOCI measurements to be mapped onto an equivalent velocity of the MS-CGS reference sample.

### 2.2 Measurement of the contrast–velocity curve and definition of the maximal static contrast

The calibration procedure relies on establishing a bijective relationship between the linear velocity of the MS-CGS sample and the *in vivo* temporal speckle contrast decrease measured by LSOCI. However, as demonstrated experimentally,^**?**^ the temporal contrast of a dynamic speckle field is not strictly bijective over the entire range of velocities. Instead, it exhibits two distinct regimes:

- **When the decorrelation time of the speckle field is much larger than the camera integration time**, the backscattered field does not evolve sufficiently during a single exposure to ensure full decorrelation between successive frames. Two acquisitions therefore remain partially correlated, and the measurement falls into a non-ergodic regime. In this limit, the temporal contrast starts from a very low value (in the absence of noise) and increases sharply as the decorrelation time decreases, approaching the value of the static spatial contrast.
- **When the decorrelation time becomes shorter than the camera integration time**, the field decorrelates fully within a single exposure. Successive measurements can then be considered statistically independent. In this ergodic regime, temporal averaging becomes equivalent to spatial averaging, and the temporal speckle contrast becomes a stable, reproducible, and monotonically decreasing function of the amount of dynamical decorrelation. This regime is the one exploited by the MS-CGS calibration and more generally by all DSI devices.

In practice, the transition from the non-ergodic to the ergodic regime occurs over a very narrow velocity range and appears as a sharp initial rise of the temporal contrast. In the present work, all practical measurements used for the MS-CGS calibration lie well within the monotonic (ergodic) regime. Consequently, the only point requiring special treatment is the temporal contrast value at the onset of the ergodic regime, where this quantity reaches its maximum. We consider here that this value corresponds to the spatial contrast measured on the static reference sample, denoted 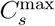, which serves as the upper bound of the calibration curve. This use of ergodicity at the very beginning of the ergodic regime will be naturally substantiated later on from the calibration curve. Practically, to measure this contrast–velocity curve, the MS-CGS sample is mounted on a directdrive rotation stage (DDR25, Thorlabs). Small Regions of Interest (ROIs) are extracted at increasing radial distances from the rotation axis. Provided they are small and sufficiently far from the center, each ROI corresponds to a well-defined linear velocity.

For each ROI, the dependence of the temporal contrast *C*_*t*_ as a function of the velocity *v* is calculated from experimental data, and the contrast loss is defined as:

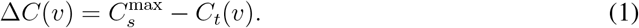

Because the MS-CGS sample satisfies all statistical requirements, the mapping Δ*C*(*v*) is smooth, strictly increasing, and highly reproducible for a given instrument. This contrast–velocity curve is the core element of the MS-CGS calibration.

### 2.3 Interpolation and inversion of the contrast–velocity relation

Once the empirical curve Δ*C*(*v*) is measured, it must be **interpolated** to allow inversion: given an observed contrast loss *in vivo*, we must estimate the *equivalent velocity* of the reference sample.

Crucially, **we do not impose any parametric model for the curve** *C*(*v*) **itself**. The literature on dynamic speckle contains numerous analytical formulations aiming to refine, for example, the Siegert relation, the temporal form of the field autocorrelation *g*^(1)^(*τ*), or the corrections needed to handle non-ergodic regimes. These models can differ substantially at low and intermediate velocities, precisely where assumptions about scattering statistics or tissue dynamics are the least robust. Yet, despite their differences, all these formulations share two structural invariants: the use of a static contrast value *C*_*s*_ and an asymptotic decay in the ergodic regime where the contrast decreases proportionally to 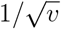.

In the MS-CGS calibration framework, we introduce a paradigm shift: the sole assumption we rely on is the equivalence of the statistical scattering properties of RBCs and the reference sample, in terms of both intensity and polarization-state distributions. Thus, we deliberately discard all model-dependent features of *C*(*v*)—no prescribed analytical form, no imposed decorrelation model. The interpolated calibration curve is therefore obtained either through a non-parametric interpolant—such as a Piecewise Cubic Hermite Interpolating Polynomial (PCHIP), which preserves monotonicity and avoids spurious oscillations—or through a minimal parametric model whose sole purpose is to accurately fit the experimental data while enabling a straightforward inversion later on. Thus, the following three-parameter expression was found to provide a good fit to our experimental data over the entire velocity range investigated (0 to 600 mm/s):

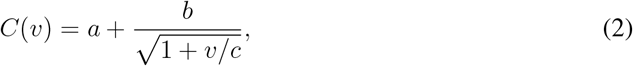

or, in terms of contrast decrease,

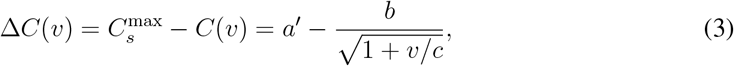

with 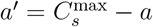.

Thus, to retrieve the equivalent velocity from a measured contrast decrease, the fitted three–parameter equation is analytically inverted as :

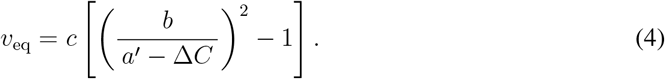

This expression allows direct computation of the equivalent velocity associated with any measured contrast decrease 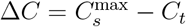, using the fitted parameters {*a*′, *b, c*}.

## 3 Experimental calibration with different laser spectral widths

### 3.1 Materials and Methods

We performed three calibrations using the following lasers: a CL-2000 diode-pumped laser from Crystal Laser with a spectral width of 0.001 nm (LASER 1), a Lambda Beam Laser PB 785-100 WL (wavelock-stabilized) from RGB Lasersystems with a spectral width of 0.1 nm (LASER 2), and a Lambda Mini Laser from RGB Lasersystems with a spectral width of 0.44 nm (LASER 3). The CMOS camera used was an aca1440-220um from Basler, combined with an HF25HA lens from FUJINON. Both LASER 2 and LASER 3 were adjusted to emit vertically polarized light. LASER 1, initially unpolarized, was filtered using a vertical polarizer. As shown in Fig. 1(a), all lasers illuminated an engineered diffuser (ED1-C20 from Thorlabs) before reaching the reference sample, ensuring homogeneous illumination. Diverging lenses (and an additional cylindrical lens for LASER 3) were used to achieve the appropriate beam size on the diffuser. The optical aperture was kept constant and optimized to obtain approximately two pixels per speckle grain. The optical powers of the three lasers were adjusted so that the same mean detected digital count, around 30 (on a scale of 0–255),^14^ was obtained after scattering on the static reference sample. A horizontal polarizer was placed just before the camera lens to implement the standard orthogonal illumination/detection configuration of the LSOCI method. For each of the six rotational speeds used to cover the full range from 0 to 600 mm/s, we computed the temporal contrast in the regions of interest (ROIs) shown in Fig. 1(b). These ROIs, positioned at increasing radii from the rotation axis, correspond to increasing linear velocities of the rotating reference sample. For each ROI, the mean temporal contrast was computed.

**Fig 1.**
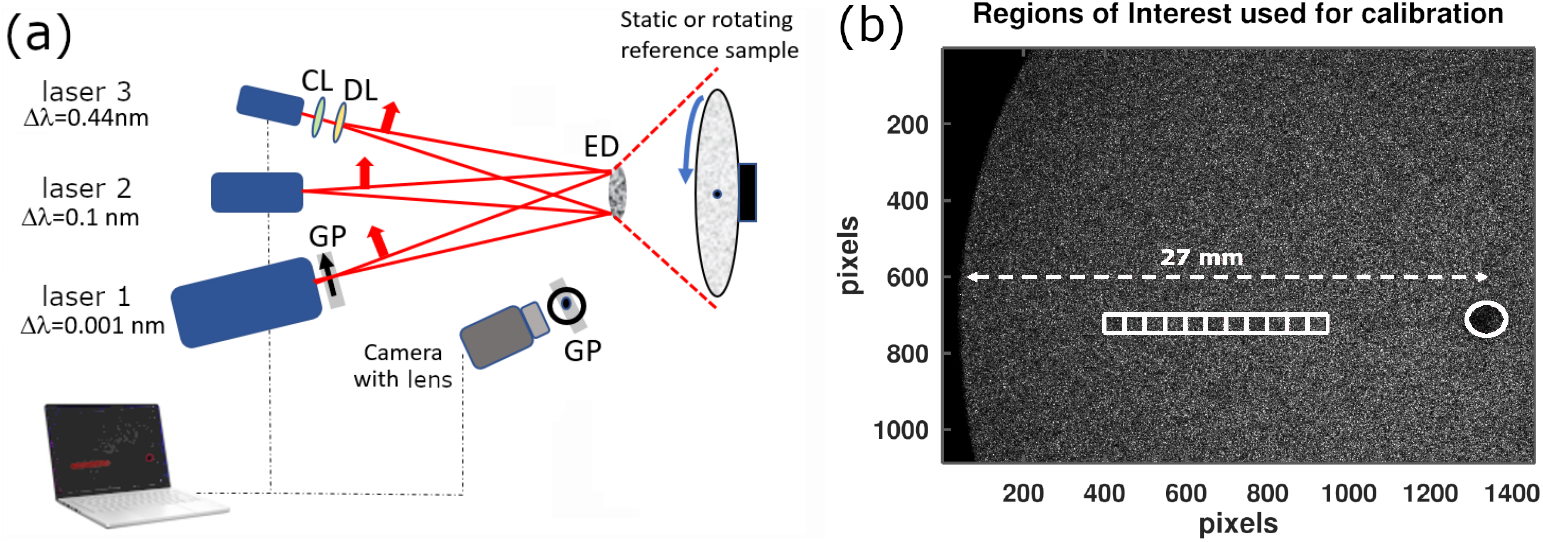
(a) Experimental setup used for the MS-CGS calibration method with three lasers having different spectral widths. Multiple rotation speeds are applied while the lasers sequentially illuminate the reference sample through an engineered diffuser. CL: cylindrical lens; DL: diverging lens; GP: grid polarizer; ED: engineered diffuser. (b) Definition of the square regions of interest (ROIs) in white on an example raw data image. The rotation axis is indicated by a white circle. Six rotation speeds are used; for each speed, several linear velocities are obtained depending on the distance of the ROI from the rotation axis.

### 3.2 Calibration results

First, when the MS-CGS reference sample was static, the maximal spatial contrast values 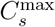 were measured as 0.8824±0.0014, 0.8870±0.0012, and 0.7157±0.0027 for laser spectral widths of 0.001, 0.1, and 0.44 nm, respectively. The uncertainties correspond to the standard deviation across the ROIs.

Second, Fig. 2 shows the decrease in contrast from its maximal value, quantified as

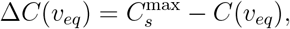

**Fig 2.**
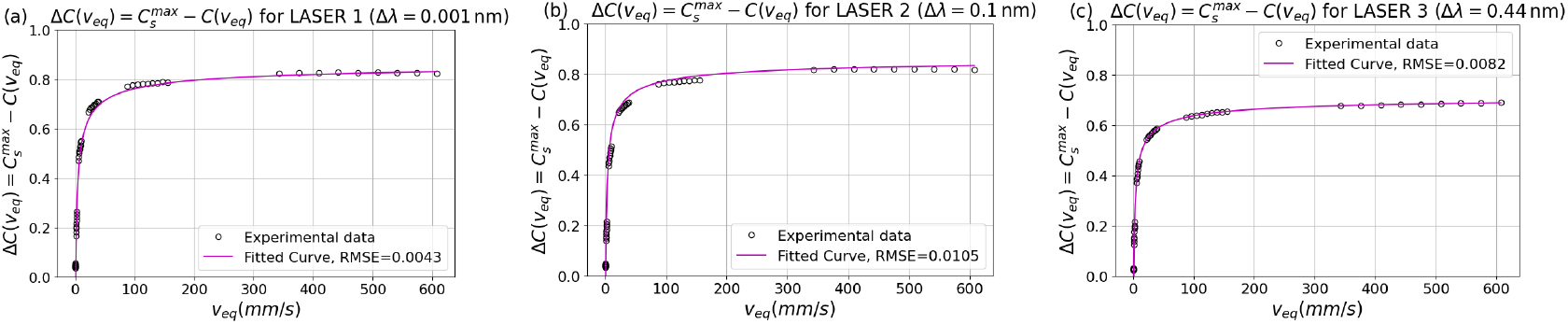
Temporal contrast decrease as a function of the linear velocity of the MS-CGS reference sample for laser spectral widths of 0.001, 0.1 and 0.44 nm in (a), (b) and (c), respectively. The CMOS camera integration time was 10 ms and the optical aperture remained constant.

for the three lasers, as a function of the linear velocity of the MS-CGS sample.

Table 1 reports the calibration coefficients obtained by fitting the experimental data using Eq. 3. The fitted values of *a* show excellent agreement with the independently measured 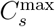 for each laser, thereby validating the previous use of ergodicity when relating the maximal temporal contrast obtainable by the instrument to the spatial contrast measured on the reference sample.

**Table 1.** Calibration coefficients for the three lasers.

|  | LASER 1 | LASER 2 | LASER 3 |
| --- | --- | --- | --- |
| | $\Delta\lambda = 0.001 \text{ nm}$ | $\Delta\lambda = 0.1 \text{ nm}$ | $\Delta\lambda = 0.44 \text{ nm}$ |
| $a$ | 0.878 | 0.877 | 0.726 |
| $b$ | 1.208 | 1.316 | 1.065 |
| $c$ | 0.929 | 0.615 | 0.691 |

Two main observations emerge from Fig. 2. First, the calibration curves for the 0.001 nm and 0.1 nm laser linewidths are very similar. This indicates that decreasing the spectral width below approximately 0.1 nm does not further improve the achievable contrast range nor significantly modify the contrast–velocity relationship. Second, the laser with the largest spectral width (0.44 nm) exhibits a markedly lower 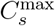, leading to a reduced dynamic range of detectable velocities. Shotnoise corrections were evaluated and found negligible for contrasts above *C* ≈ 0.1. All three lasers were operated to produce speckle patterns with mean digital counts around 30, ensuring operation in the linear regime of the CMOS detector. At this signal level, dark noise can also be neglected. As an illustration, a contrast decrease of Δ*C* = 0.6 obtained with the 0.44 nm laser linewidth corresponds, via Eq. 4, to an equivalent velocity *v*_*eq*_ ≈ 50 mm/s. The same biological region imaged with the 0.1 nm or 0.001 nm laser linewidths would exhibit slightly different raw contrast decreases (around 0.7), but, thanks to the calibration, would yield the *same* equivalent velocity of 50 mm/s.

### 3.3 In vivo measurement comparison between two calibrated instruments

Using the LSOCI setup (orthogonal illumination/detection protocol) with LASER 1 and LASER 3, we performed dynamic speckle imaging of a minor finger wound. This model was chosen because localized skin injuries are known to induce a strong and spatially heterogeneous microcirculatory response in the perilesional region, making it an ideal test case for evaluating calibration consistency. The resulting perfusion dynamics naturally spans healthy skin, transitional reactive tissue, and the damaged zone.

Figure 3 shows the temporal speckle contrast obtained with (a) LASER 1 (Δ*λ* = 0.001 nm) and (b) LASER 3 (Δ*λ* = 0.44 nm). Although both images exhibit very similar spatial patterns, their contrast levels differ substantially, as expected from the strong dependence of 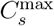 on the laser spectral width.

**Fig 3.**
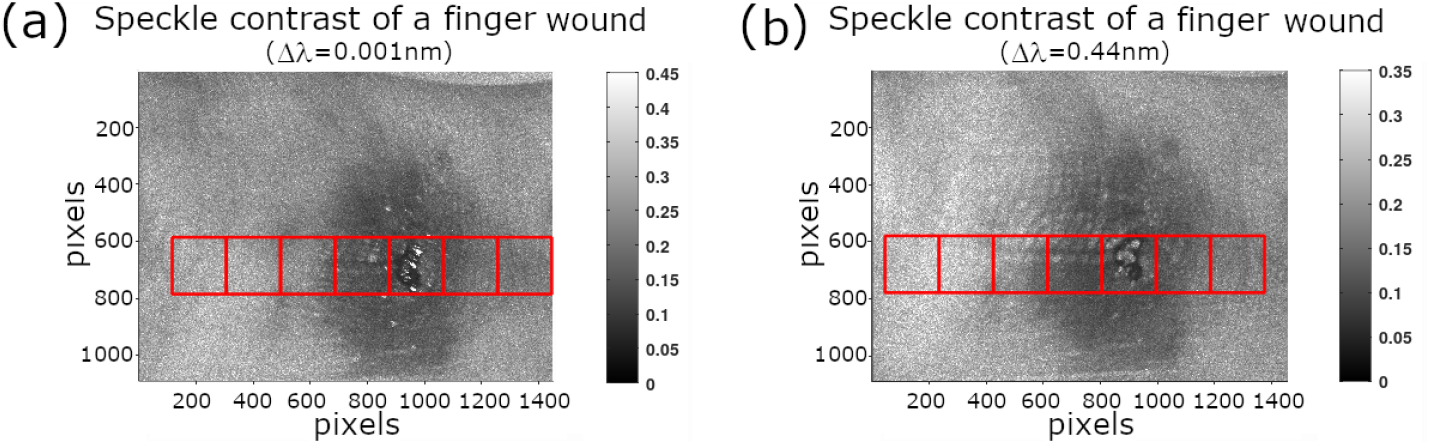
Temporal speckle contrast of a finger wound obtained using laser spectral widths of (a) 0.001 nm and (b) 0.44 nm. Despite the visual similarity of the two images, the absolute contrast values differ significantly because of the spectral-width dependence of 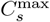.

Seven square ROIs (indicated in Fig. 3) were positioned along a transect crossing healthy skin, the perilesional zone, and the wound center. The mean temporal contrast in each ROI is reported in Fig. 4(a). Without calibration, the two lasers yield clearly different contrast values. Using the proposed MS-CGS calibration, the contrast decrease relative to each instrument-specific 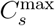 is mapped bijectively to the equivalent velocity *v*_*eq*_ of the reference sample (as derived from the calibration curves in Fig. 2). This enables a direct comparison of microcirculatory activity across instruments.

**Fig 4.**
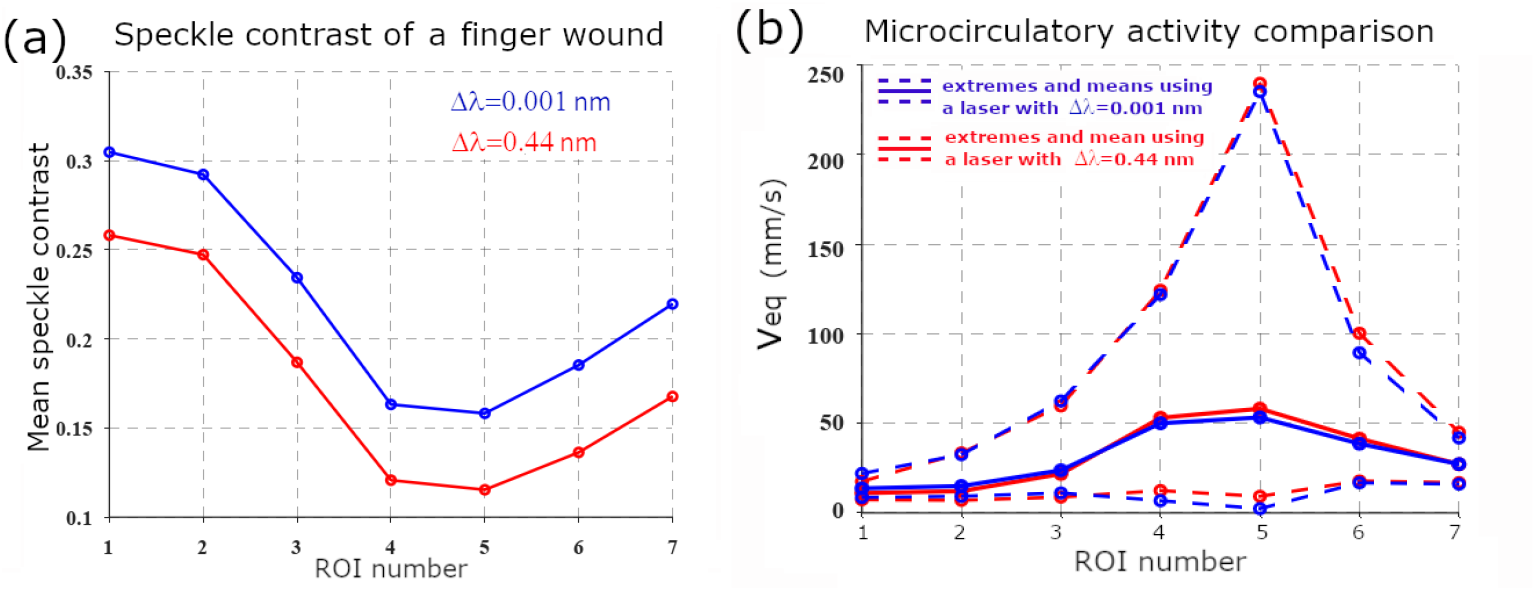
(a) Mean temporal speckle contrast within the ROIs defined in Fig. 3, obtained with LASER 1 and LASER 3. (b) Corresponding equivalent velocity *v*_*eq*_ after MS-CGS calibration. Although raw contrasts differ strongly, the calibrated equivalent velocities exhibit excellent agreement across all ROIs.

Figure 4(b) shows the corresponding *v*_*eq*_ values for the same ROIs. After calibration, the two lasers exhibit excellent agreement across the entire range of microcirculatory activity, despite (i) the sequential acquisition of the two datasets, which unavoidably introduces biological variability, and (ii) slight finger motion relative to the optical head during the acquisition. This latter bias was minimized by using relatively large ROIs; however, it remains a known limitation for pixel-level flow quantification. In previous work, we addressed this issue by implementing LSOCI in contact mode,^5, 14^ which significantly improves spatial stability.

Finally, Fig. 5 provides a global comparison of the equivalent velocity maps obtained with both lasers using a common color scale. The two calibrated images exhibit nearly identical spatial distributions and magnitudes of microcirculatory activity, demonstrating the robustness of the MS-CGS calibration method for inter-instrument comparison.

**Fig 5.**
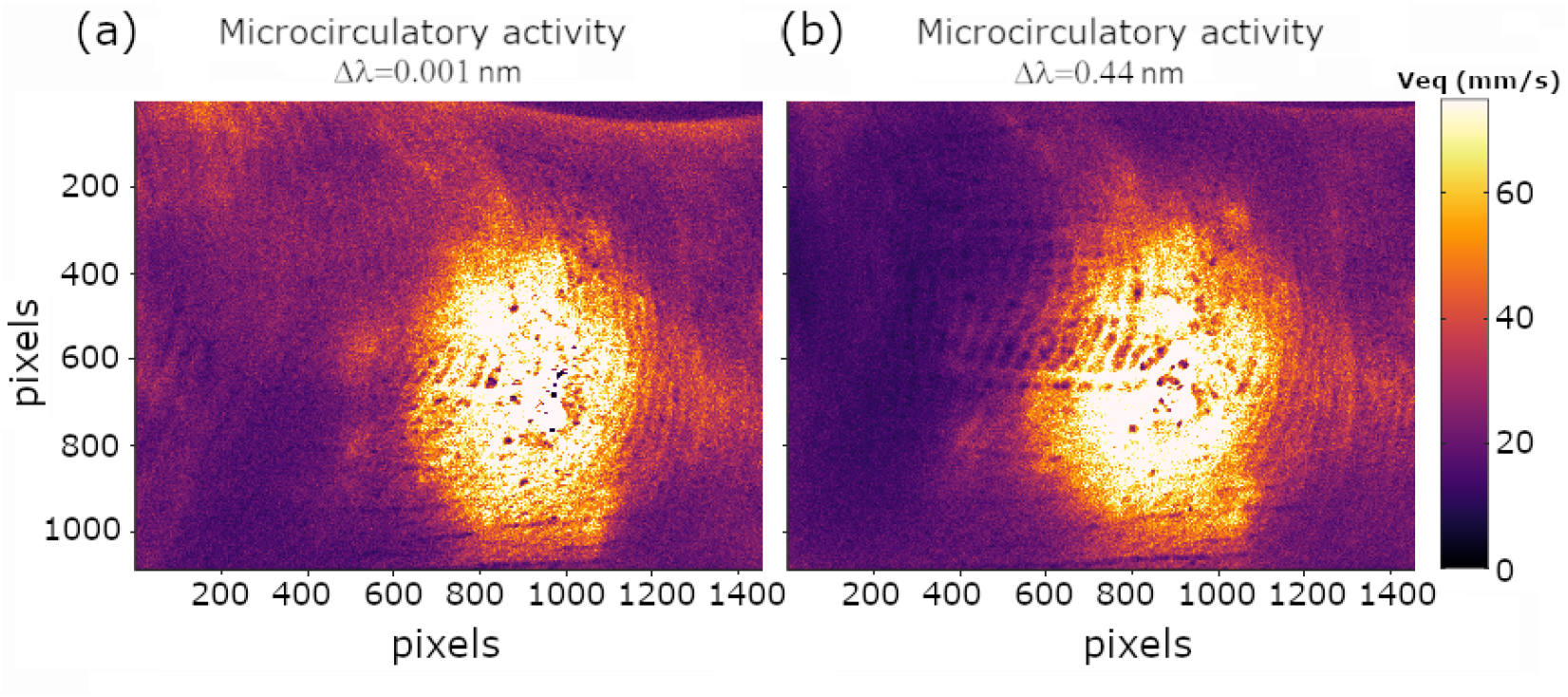
Equivalent-velocity maps (mm/s) obtained after MS-CGS calibration using the two lasers with different spectral widths. Both images use the same color scale and reveal nearly identical microcirculatory patterns across the entire field of view.

## 4 Discussion

The proposed MS-CGS calibration relies solely on the assumption that multiple scattering within and between moving RBCs is of sufficiently high order to generate enough entropy in the backscattered polarimetric states to uniformly cover the surface of the Poincaré sphere. When this assumption is fulfilled, moving RBCs generate circular Gaussian speckle patterns that are fully spatially depolarized. This scattering can then be compared with that of a moving reference sample, which produces dynamic speckle patterns with similar statistical properties in both intensity and polarization states distribution. A decrease in speckle contrast for each pixel during *in vivo* imaging can therefore be directly related to the speed of such a reference sample. It is essential to emphasize that the quantity retrieved through this calibration is not the absolute velocity of individual RBCs, nor any specific directional component of flow. Instead, the equivalent velocity *v*_eq_ represents a macroscopic measure of the overall three-dimensional agitation of RBCs within the elementary volume where multiple scattering occurs. This interpretation is particularly well suited to the dense, highly interconnected 3D architecture of the microvascular network, where multiple scattering is inherently strong.

The experimental results lead to two major conclusions. First, no significant differences were observed between lasers with spectral widths below 0.1 nm. The maximum speckle contrast 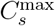 does not increase further when narrowing the linewidth from 0.1 nm to 0.001 nm, and the calibrated contrast–velocity curves remain similar. This observation provides a practical guideline for selecting laser sources for LSOCI: linewidths narrower than 0.1 nm offer little additional performance benefit.

Second, despite their very different laser spectral widths (0.001 nm and 0.44 nm), two MS-CGS– calibrated LSOCI systems produced highly consistent equivalent-velocity measurements *in vivo* across a dynamic range extending from a few mm/s up to about 250 mm/s. This agreement strongly supports the validity of the MS-CGS approach for inter-instrument standardization.

However, some bias may be expected if the order of multiple scattering is too low and therefore fails to uniformly cover the surface of the Poincaré sphere. So far, our various observations on human skin have always fulfilled the required assumption when working in the near-infrared domain. A change of wavelength, however, could compromise the method by reducing the order of scattering. Moreover, classical LSCI, without an orthogonal illumination/detection configuration, retains the polarized surface contribution of the backscattered speckle and is therefore also expected to violate the MS-CGS hypothesis due to the resulting inhomogeneous coverage of the Poincaré sphere.

Overall, the MS-CGS calibration addresses a longstanding gap in dynamic speckle imaging: it provides a simple, rapid, and robust means of producing quantitatively comparable measurements across different LSOCI instruments. By grounding the calibration on a multiple-scattering reference rather than on a physical model of tissue dynamics or sensor design, it offers a pragmatic and reliable route toward standardized microcirculation quantification.

## Conclusion and perspectives

We have introduced and validated a calibration method for LSOCI that enables robust and quantitative assessment of microcirculatory activity. The key contribution of this work is the definition of an *equivalent velocity*, derived from a multiple-scattering reference sample whose statistical properties are fully controlled. By comparing the contrast decrease observed *in vivo* to that produced by the moving reference sample, LSOCI measurements can be expressed in terms of a single, interpretable quantity that is independent of the instrument and of any detailed model of tissue dynamics. This equivalent velocity does not represent the absolute motion of individual red blood cells, but rather a macroscopic measure of their three-dimensional agitation within the multiply scattering volume. Such a metric is naturally suited to the dense, heterogeneous, and volumetric architecture of the microvascular network, where isolating directional flow components is neither feasible nor necessary for most biomedical applications. Once implemented, the MS-CGS calibration can be performed within seconds, and it provides a unified scale on which different LSOCI instruments—despite variations in laser linewidth, optical aperture, or detector characteristics—can be directly compared. This capability opens the way to standardized quantification of microcirculation across instruments and clinical sites. It also enables longitudinal monitoring of pathological progression or therapeutic response, without requiring a stable hardware configuration over time. Given the compactness and robustness of LSOCI, together with the simplicity of the proposed calibration procedure, we believe that the MS-CGS framework provides a practical foundation for the widespread adoption of quantitative dynamic speckle imaging in both research and clinical environments.

## Author Contributions

**Xavier Orlik** received master’s degrees in electromagnetism (INPT, 1997) and medical imaging (UPS, 2008), and a Ph.D. in 2001 within the European program *Near-Field Optics for Nanotechnology*. He joined ONERA in 2003, where he developed research on dynamic speckle, polarimetric imaging, and random electromagnetic fields. He obtained his Habilitation à Diriger des Recherches in 2012 and has been Research Director since 2022. His work focuses on non-conventional optical imaging, with applications ranging from surface characterization to biomedical optics. He is the inventor of four international patents and teaches optical imaging and electromagnetism at É cole Polytechnique and ISAE-SUPAERO.

**Aurélien Plyer** graduated from Université Pierre et Marie Curie (Paris 6) in 2008 and received his Ph.D. in image processing from Université Paris 13 in 2013. He is a researcher at ONERA, the French Aerospace Lab, within the Information Processing and Systems Department. Since 2024, he has been heading the research unit *MIC — Measurement, Image, Codesign*. His work focuses on high-efficiency parallel algorithms for low-level video processing, 3D environment perception, and large-scale data fusion, with applications spanning experimental physics, robotics, and remote sensing.

**Elise Colin** received the Dipl. Ing. in electrical engineering from Supelec and the M.Sc. degree in theoretical physics from the University of Paris XI, France, in 2002. She obtained her Ph.D. from the University of Paris VI in 2005, followed by the *Habilitation à Diriger des Recherches* from the University Paris-Sud, Orsay, in 2014. She has been a Research Director at ONERA since 2020. Her research focuses on advanced and non-conventional imaging modalities, with a particular emphasis on polarimetry across radar and optical regimes, electromagnetic wave scattering, and the development of innovative sensing strategies for complex media.

